# Decomprolute: A benchmarking platform designed for multiomics-based tumor deconvolution

**DOI:** 10.1101/2023.01.05.522902

**Authors:** Song Feng, Anna Calinawan, Pietro Pugliese, Pei Wang, Michele Ceccarelli, Francesca Petralia, Sara JC Gosline

**Author notes:** Correspondence should be addressed to Sara Gosline.

## Abstract

Tumor deconvolution is a reliable way to disentangle the diverse cell types that comprise solid tumors. To date, however, both the algorithms developed to deconvolve tumor samples, and the gold standard datasets used to assess the algorithms are geared toward the analysis of gene expression (e.g., RNA-seq) rather than protein levels in tumor cells. While gene expression is less expensive to measure, protein levels provide a more accurate view of immune markers. To facilitate the development as well as improve the reproducibility and reusability of multi-omic deconvolution algorithms, we introduce Decomprolute, a Common Workflow Language framework that leverages containerization to compare tumor deconvolution algorithms across multiomic data sets. Decomprolute incorporates the large-scale multiomic data sets produced by the Clinical Proteomic Tumor Analysis Consortium (CPTAC), which include matched mRNA expression and proteomic data from thousands of tumors across multiple cancer types to build a fully open-source, containerized proteogenomic tumor deconvolution benchmarking platform. The platform consists of modular architecture and it comes with well-defined input and output formats at each module. As a result, it is robust and extendable easily with additional algorithms or analyses. The platform is available for access and use at http://pnnl-compbio.github.io/decomprolute.

**Motivation:** To provide a comprehensive platform for algorithm developers and researchers to benchmark and run tumor deconvolution algorithms on multiomic data.

## Introduction

Tumor growth and metastasis rely on the exchanges between the tumor cells and additional components constituting the tumor microenvironment^1^. Understanding the interactions between the tumor cells and surrounding non-malignant cells, including stromal, endothelial and immune cells, is essential to model the mechanisms underlying tumor survival and spreading^2^. In particular, identifying the degree and nature of immune cell infiltration can assist in predicting a tumor responsiveness to specific immunotherapeutic regimens^3,4^. Hence, new technologies, such as mass cytometry^5−8^ and single-cell RNA sequencing^9−12^, have been applied also to study tumor microenvironment together with *ad hoc* computational algorithms to deconvolve cell types from bulk molecular measurements^13−17^.

Algorithmic tumor deconvolution is based upon the knowledge that specific genes are expressed at distinct levels within specific cell types^18^. Given the prior knowledge of the specific combinations of gene expression levels to expect in a specific cell type, numerous existing computational algorithms can provide estimates of relative cell types present in the profiled tissue. One such algorithm, Microenvironment Cell Populations-counter (MCP-counter), provides tumor deconvolution predictions from bulk RNA sequencing data using a gene signature matrix. This signature matrix is derived from previously published gene expression datasets, which are analyzed to provide an MCP-counter score, comprised of the logarithm of geometric mean of the marker genes for each cell type^17^. CIBERSORT and CIBER-SORTx employ a linear modeling approach to estimate celltype composition from a signature matrix and bulk gene expression matrix ^14,19^. EPIC (Estimating the Proportions of Immune and Cancer cells) is an algorithm that uses a similar approach to CIBERSORT but normalizes the gene expression values to account for proportions of healthy vs. malignant cells ^16,20^. xCell expands upon these existing numeric approaches by leveraging gene set enrichment statistics to characterize cell types^15^. These methods, in addition to many others not explicitly mentioned here^21^, showcase the great need for tumor deconvolution from bulk measurements.

While tumor deconvolution algorithms are highly effective at using gene expression data, their performances are unexplored on proteomic data despite the rise ofcancerspecific proteomic datasets^22−25^ together with the established fact that protein levels do not always correlate with mRNA ^26−29^, suggests that algorithmic deconvolution could be more effective if a proteomic signature matrix or proteomics-derived signatures are utilized. Recent work by Rieckmann et al.^30^ has created a dataset that enables, the definition of immune cell types based on proteomic data.. However, there is still no available ‘gold standard’ dataset to evaluate the ability of an algorithm using these proteomics-defined immune cell types to deconvolve tumor data. On the other side, for mRNA-based deconvolution, there are numerous single-cell datasets as well as sorted cell experiments that can be used for such benchmarks^31−33^ which are missing for proteomic data.

Here we introduce Decomprolute, a containerized set of scientific workflows that enables the community to compare the performance of existing or novel deconvolution algorithms specifically on proteomic data. We demonstrate the utility of Decomprolute using a subset of published deconvolution algorithms on the CPTAC3 cancer datasets (cite data resource paper) for direct comparison with mRNA-based algorithms, simulated data, and pan-cancer immune subtypes. The framework comprises four existing deconvolution algorithms but can easily work with any new algorithm that is able to function in a Docker container. Our system is flexible enough to accept additional signature matrices, deconvolution algorithms and datasets, both as input and for validation, as we hope that it will inspire future development in the tumor deconvolution space.

## Results

### Modular workflow framework enables flexible comparison of deconvolution results across signatures, cancer types and algorithms

The goal of Decomprolute is to encourage rapid development and benchmarking of novel deconvolution algorithms and cell type signature matrices. As such, the underlying architecture is structured around a modular framework that allows additional algorithms, datasets or cell type signatures to be easily added for comparison. The platform enables users to run and generate figures for experiments in a reproducible fashion and its modularity allows it to be expanded to run additional statistical tests as needed.

The overall software architecture is described in **Figure 1**. The modules that comprise decomprolute are: 1) **prot-data**, which accesses data from published cancer proteomics resources (cite pan can resource paper), 2) **mrnadata**, which accesses matched gene expression data from the same patients (cite same), 3) **signature-matrices**, which returns specific signature matrices to evaluate with existing algorithms^15−17,19^, 4) **tumor-deconv-algs**, which evaluates a combination of gene and/or protein expression data and signature matrix on an algorithm of interest, and 5) **metrics**, which compares the performance of the algorithm on a set of benchmarks we define. Each module takes a standard set of inputs and outputs and therefore can be interchanged and appended. A full list of parameters is described in **Table 1**. This modular design enables users to plug in their own data or algorithms or create a new evaluation metric by which they can compare data.

**Table 1:** Overview of Decomprolute modular arguments and description.

| Module | Inputs | Outputs |
| --- | --- | --- |
| prot-data | Cancer Type (e.g., LUAD)<br>Tissue type (e.g., tumor) | A single file of protein expression across patients |
| mrna-data | Cancer type (e.g., LUAD)<br>Tissue type (e.g., tumor) | A single file of gene expression across patients |
| signature-matrices | Signature name | A single file of transcriptomic or proteomic profile chosen for cell types |
| tumor-deconv-alg | Cancer type<br>Signature name<br>Algorithm name | A single matrix where rows are cell types, samples are columns, and each value is the estimated fraction of that cell type for that sample |
| metrics | Specific parameters based on the type of metrics | Figures and tables summarizing cross-algorithm analysis |

**Figure 1:**
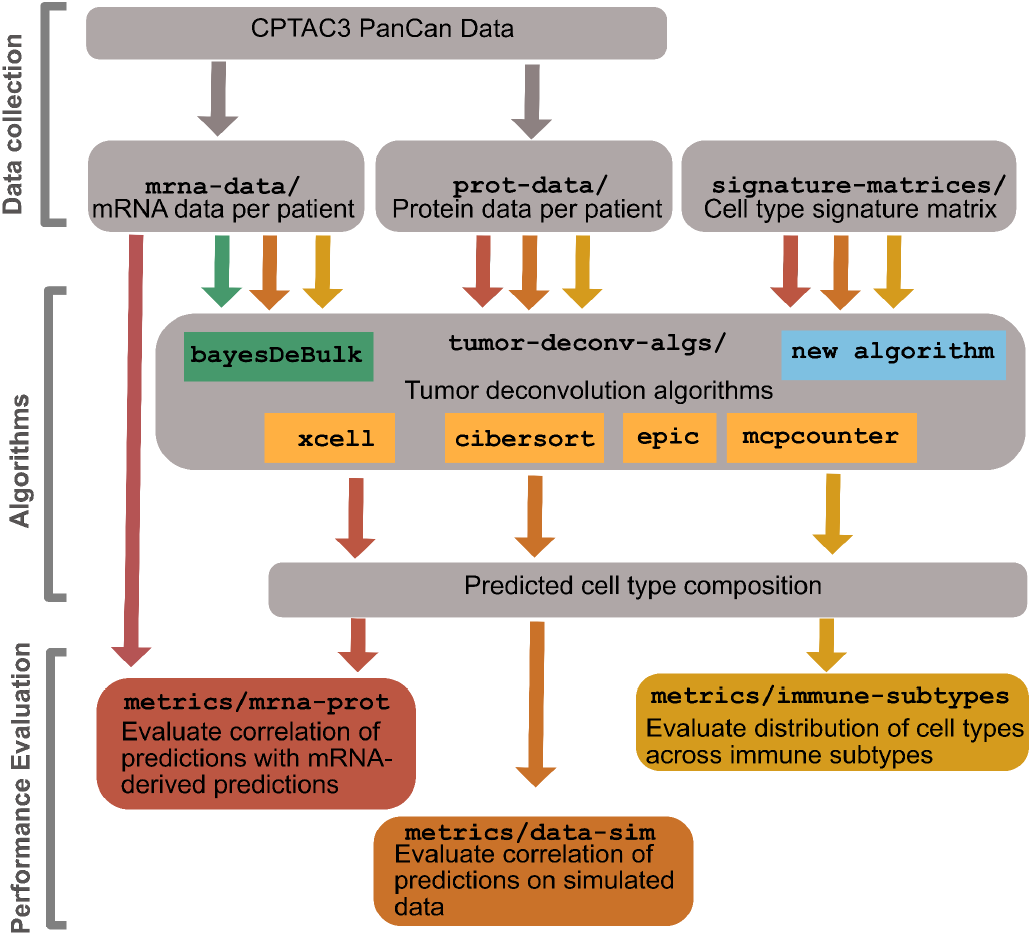
Overview of Decomprolute modular architecture describes four primary subdirectories of Decomprolute. The mrnadata and prot-data modules both pull from the CPTAC pan-cancer resource. The signature-matrix module contains genes that represent different cell types, the tumor-deconv-algs module contains each of the algorithms we implemented, while the metrics module contains all the modules used to measure performance.

While designed for flexible extensibility, the platform includes built-in data access scripts, signature matrices, and deconvolution algorithms. We focused on publicly available CPTAC data across 10 cancer cohorts (**Table 2**) (cite data resource paper) that consist of matched proteomic and transcriptomic measurements. This allows the user to evaluate performance across different tumor types. We also implemented four publicly available algorithms for deconvolution to provide examples of how these can be used and compared in practice. We include signature matrices that were published from mRNA expression profiles^14,19^ and generated new ones from sorted proteomic data ^30^, as described in the Methods. Lastly, we developed three metrics that enable users to compare and contrast various aspects of tumor deconvolution on proteomics data: 1) evaluation on simulated data, 2) agreement between mRNA and protein, and 3) comparison with immune subtypes ^34^. Each of these are demonstrated below.

**Table 2:** Summary of samples by cancer type.

| Cancer type | Tumor samples | Normal samples |
| --- | --- | --- |
| BRCA | 122 | 0 |
| CCRCC | 103 | 80 |
| COAD | 110 | 100 |
| GBM | 99 | 0 |
| HNSCC | 108 | 62 |
| LSCC | 108 | 99 |
| LUAD | 110 | 101 |
| OV | 83 | 20 |
| PDAC | 105 | 44 |
| UCEC | 95 | 18 |

### Simulated data metric enables determination of proteomics-derived signature matrix

To provide a data independent assessment of deconvolution algorithms, we first developed a suite of simulated datasets that allow each algorithm to be compared in an unbiased fashion. Specifically, we simulated a dataset from proteomic data as well as from mRNA data as described in STAR methods. We were then able to compare, for both proteomic and mRNA datasets, how well different cell types were predicted by correlating these values with ground truth on simulated data. This test can be run using the metrics module available on our website.

To showcase the value of such a metric, we compared two proteomic signature matrices to determine how well different algorithms can deconvolve simulated samples generated using the proteomic immune data from Rieckmann et al.^30^. The signature matrices were also derived from this dataset (see Methods). Briefly, from the original 26 sorted immune cell types, our original analysis found that many of them were quite similar in proteomic profile, suggesting that the proteins used would not be specific enough to distinguish different cell types. As a result, we grouped the 26 cell types into 9 different categories to create a new LM9 signature matrix. Using the LM9 signature matrix to deconvolve the simulated data provides heterogenous results across algorithms, as depicted in **Figure 2A**. The correlation with simulated levels of CD4 T cells, for example, is low across most algorithms but CIBERSORT (yellow). We then compared this approach to what we called the LM7c signature matrix, which starts with the LM9 but compresses basophils, eosinophils and neutrophils into a single ‘granulocyte’ category (see STAR Methods). As a result, we are able to predict the immune cell types with higher overall correlation and also see improvements within cell types (**Figure 2B**), suggesting that the LM7c signature matrix is more accurate for proteomics-based deconvolution.

**Figure 2:**
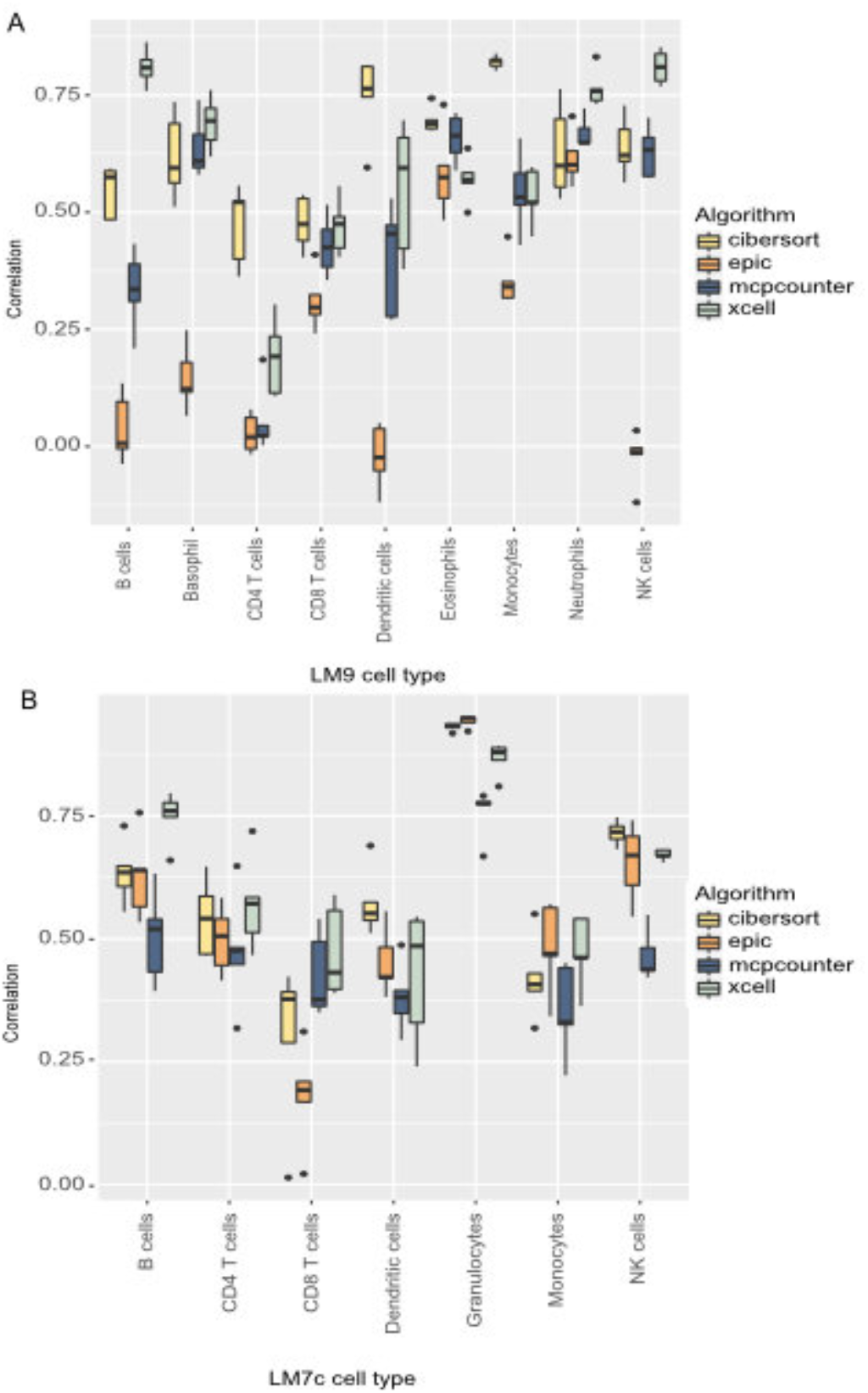
Correlation of proteomics-derived matrices. Correlations are between predicted results and simulated results from the LM9 and (B) LM7c signature matrices. X-axis depicts predicted cell type and y-axis is spearman rank correlation across 10 simulated datasets between algorithm prediction (color indicated on right) and known fractions of cells.

### Assessing algorithmic agreement between protein and mRNA-based deconvolution

As we described earlier, mRNA-based tumor deconvolution algorithms^14−17^ have demonstrated success when compared to gold standards. These datasets contain known mixtures of individual cell types or paired single-cell measurements, alongside bulk RNA-seq data. Hence, it is possible to compare different algorithms to see which method deconvolves better having a ground truth as reference. However, because no such dataset exists for tumors measured via bulk proteomics, we use the bulk mRNA measurements matched to bulk proteomic measurements across 10 different cancer types from the CPTAC 3 pan-cancer efforts to identify which algorithm-signature matrix combination gives the best results on proteomic data when compared to mRNA-based predictions.

Like our other metrics, this test is also implemented as a single workflow that produces numerous tables and figures for further analysis. **Figure 3** depicts a subset of the results of this analysis, measuring the concordance between mRNA and protein-based deconvolution using one of two different distance metrics: the Jensen-Shannon divergence and Spearman rank correlation (see STAR Methods). Across the ten cancer types and three signature matrices, the MCP-counter algorithm showed the highest amount of agreement for its predictions on mRNA and protein data, using both transcriptomics- and proteomics-derived signature matrices, as depicted by low average distance in **Figure 3A** and high correlation in **Figure 3B**. The xCell algorithm achieves similar results, with a low distance (3A) and high correlation (3B) between mRNA and protein-based predictions. CIBERSORT and EPIC, which do not rely on gene signatures but on gene expression values, proved to be less flexible across mRNA and protein deconvolution algorithms.

**Figure 3.**
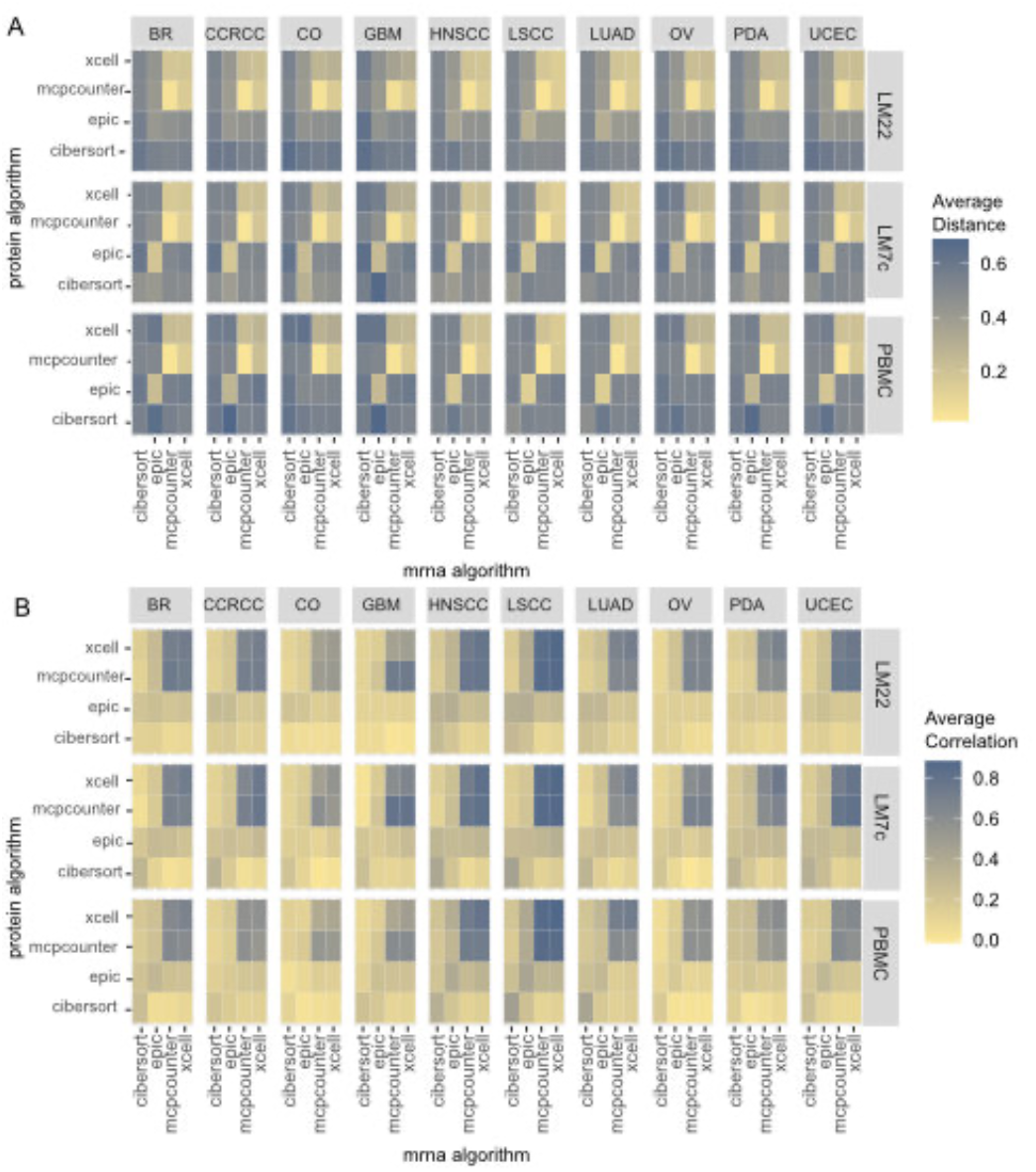
All vs. all algorithmic comparison. Average (A) Jenson-Shannon divergence distance and (b) Spearman rank correlation between deconvolution results run on mRNA data (x-axis) and proteomics data (y-axis). Individual values are divided across signature matrices and cancer types. Legend along right side.

### Cell type-specific variation in algorithmic performance

Due to biases in the algorithms and signature matrices, we also provided the ability to compare mRNA and protein deconvolution results across algorithms and cell types. Specifically, we measured, for each cell type, the correlation between mRNA and protein across patient samples. The results are depicted in **Figure 4**. Here, we see that even within the same signature matrix, we can get different degrees of correlation between mRNA and protein algorithms. As we see in **Figure 4**, MCP-counter and xCell have a high degree of agreement between mRNA (columns) and proteins (rows). However, xCell and MCP-counter have a lower degree of correlation when it comes to NK cells, T4 cells, and DC cells. Generally, the LM7c signature matrix, which is derived from protein data, has higher mean correlation than the PBMC matrix, which comes from sorted mRNA-based measurements.

**Figure 4.**
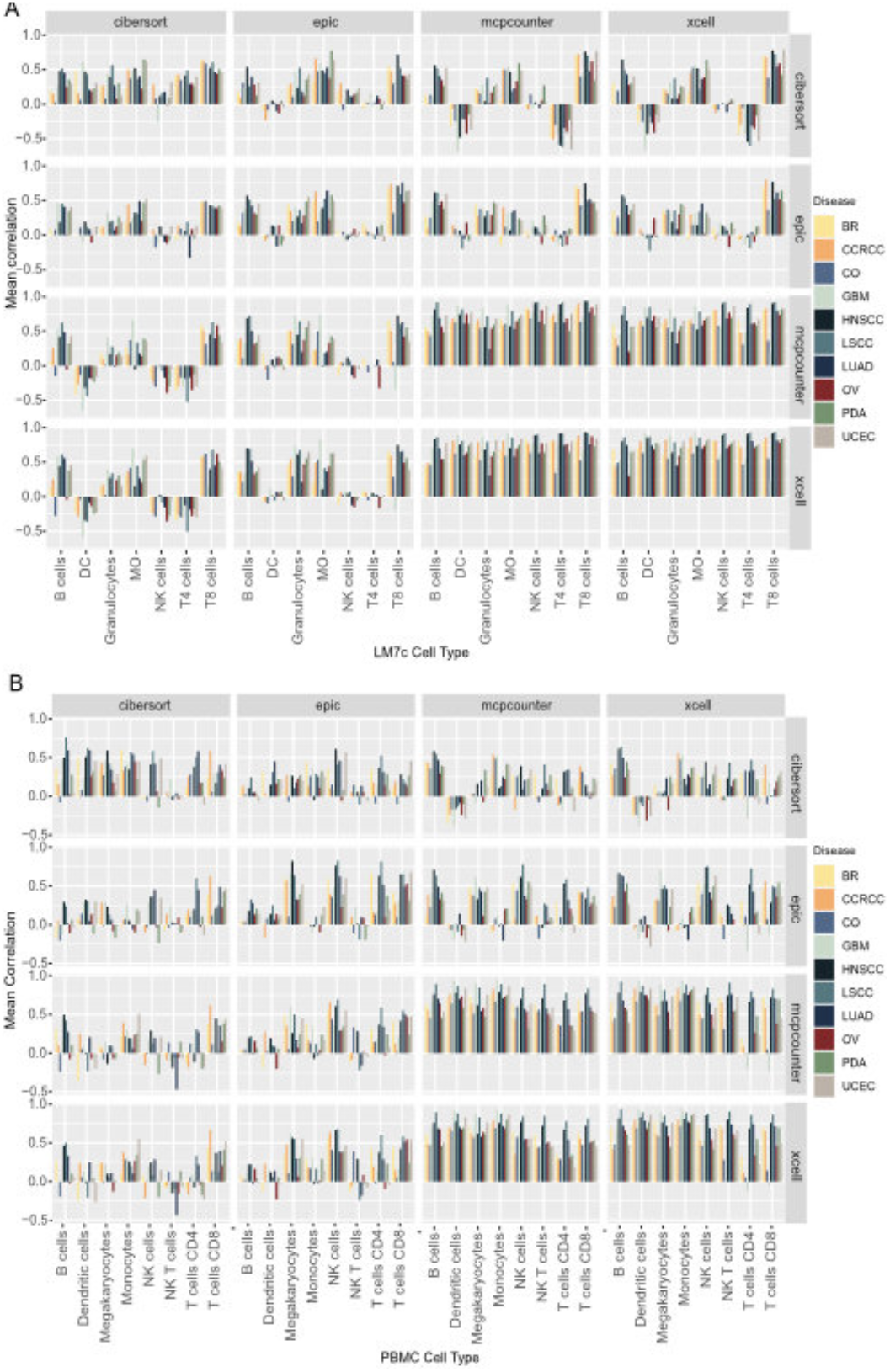
Correlation across algorithms and cancer by cell type. Predicted cell types across tumors by algorithm and cancer type for the (A) LM7c signature matrix and (B) PMBC signature matrix. Columns represent algorithms run on mRNA data and rows represent algorithms run on protein data. The correlation for each cancer type and cell type are shown bars colored by cancer type.

### xCell captures immune subtypes in proteomic-derived cell type composition

As a third validation we again leveraged CPTAC 3 proteomics datasets. Here, we utilized the classification of each tumor sample into one of the six immune subtypes^34^ predicted using the mRNA data^35^. Specifically, we compared the deconvolution results of the four algorithms on tumor samples of all 10 tumors respect to the immune subtypes classification. The values from each algorithm and for each cell type were transformed to z-scores. We used PBMC on RNA-seq data and LM7c on proteomic data and the median as summary statistics, since we are interested in how many samples have a z-score above or below zero. We focused on immune subtypes with more than one hundred samples assigned (Figure 5).

**Figure 5.**
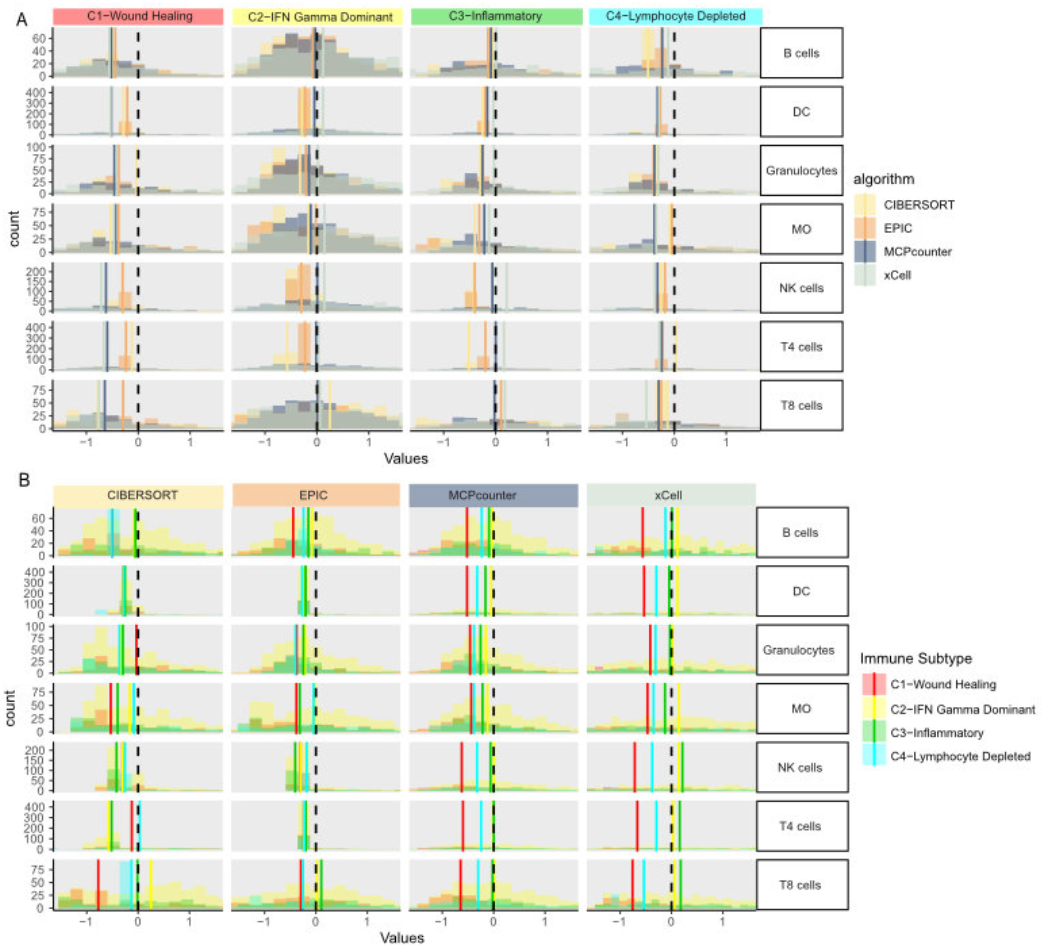
Predicted cell types across tumors by immune subtype and algorithm. (A) Depicts distribution of cell types (rows) as predicted by various algorithms. Color of density plots describes the algorithm used to score the subtypes. (B) Depicts the same distribution of values, but the columns represent algorithms and the color of the density plots represents the subtypes. Dashed bars represents a Z-score of 0, while the colored bars represent the median of each histogram.

We can use the cross section of immune signatures with cell type to evaluate how accurately the algorithms can predict immune activity. For example, the C1 (wound healing) immune subtype, characterized by a high proliferation rate, shows no enrichment for any cell type for both types of data and for all algorithms (**Figure 5A**). The C2 (IFN-g dominant) subtype, defined rich of CD8 T cells and M1 macrophages, has the highest number of samples assigned, 505. Interestingly, xCell predicts, for the samples assigned to this cluster, an enrichment for most immune cells regardless of the type of data used while the other algorithms show an opposite result (**Figure 5B**). CIBERSORT found enrichment of CD8 T cells in the C2 samples. These samples serve as a good benchmark of immune activity because they can be seen as “immune-hot” - the IFN-g response that characterizes this immune subtype, causes the activation of both innate and adaptive immune system. The C3 subtype, defined as inflammatory, shows enrichment for lymphocytes for both types of data with xCell whereas the C4 subtype, defined as lymphocytes depleted, shows a minimal enrichment for both types of data for CD4 T cells with CIBERSORT. Overall, xCell is more accurately able to capture the immune activation in this subtype compared to the other algorithms.

## Discussion

Here we introduced a benchmarking platform to assess the performance of tumor deconvolution algorithms on proteomics data. It is comprised of four modules, each of which can be altered to allow for additional (1) algorithms, (2) proteomic datasets, (3) signature matrices, and (4) evaluation metrics. We showcase each of the evaluation metrics. First, we show how the algorithms can be run on simulated data using both proteomics and mRNA expression levels. We then compare mRNA-based deconvolution to proteomics to determine how well the algorithms agree. Lastly, we use mRNA-derived immune subtypes to evaluate how proteomics-based tumor deconvolution algorithms identify relative cell fractions within each subtype.

The need for such a system emerges from the absence a protein-native tumor deconvolution ‘gold standard’ that can be used to evaluate the performance of existing tumor deconvolution algorithms on proteomic data. In the absence of such a dataset, we employ these three metrics - data simulation, correlation, and immune analysis, to enable the measurement of existing algorithms. We hope that such a platform will be used by the community to further develop tumor deconvolution algorithms based on proteomic data so that we can get more insights from the inference of cell phenotypes using this data.

As we learn more about the value of proteomics measurements in cancer studies (Cite), understanding the nuances of proteomics data in tumor deconvolution is highly valuable. This framework will facilitate the development of proteomics and proteogenomic tumor deconvolution algorithms by providing an easy way to compare newly developed approaches to those that already exist. We believe this platform is robust to additional datasets, algorithms, and signature matrices and will be broadly used by the tumor proteomics community.

## Limitations of study

We only experimented with four tumor deconvolution algorithms for which we were able to build containers. Some tools, such as CIBERSORTx ^19^, were not freely available to be made compatible with our framework and therefore could not be evaluated.

## Acknowledgements

We would like to acknowledge the CPTAC 3 PanCancer Immune working group for their valuable feedback. PNNL is operated for the DOE by Battelle Memorial Institute under Contract DE-AC05-76RL01830

## Author contributions

SF: Lead developer on Docker containers and workflows, wrote manuscript

AC: Tested scripts and supported documentation, wrote manuscript.

FP: Provided guidance, wrote manuscript

PW: Provided guidance, wrote manuscript

PP: Evaluated tumor immune, wrote manuscript.

SG: Led project development, wrote manuscript.

## Resource availability

### Materials Availability

All materials used for this analysis are published and freely available as part of the CPTAC data resource paper.

### Data and Code Availability

All source code is publicly available via Github at https://github.com/pnnlcompbio/decomprolute. In addition to the underlying software for executing and assessing the performance of the various algorithms, this repository includes signature matrix files, dummy test data, sample inputs, and the CI/CD configuration file. The CWL workflows to execute the pipeline are organized under a single directory. From this directory, users can execute deconvolution and performance comparison. If executing the code on a local machine, output is saved directly in this directory. Using a Docker image will save the output on the corresponding container directory, and the user can transfer the file to their local computer with a mounted directory or with the ‘docker scp’ command.

## Materials and Methods

### Cancer transcriptomic and proteomic data

We provide a flexible framework that enable both the mRNA and proteomics data to be handled in individual modules to make it easier to upgrade and replace these modules with updated data as additional proteomics datasets are released. Specifically, we rely on the CPTAC Python package36 in attempts to build a framework that would be flexible with respect to incoming data. Therefore, Decomprolute can be used with this package or replaced with other similar packages or data files. Table 2 lists the sample numbers available at the time of publication.

### Tumor deconvolution algorithm modules

Within the tumor-deconv-algs module we currently have implemented four distinct algorithms from the community: CIBERSORT^14^, MCP-counter^17^, xCell^15^, and EPIC^16^. Additional algorithms can be added provided they take a tab-delimited file as input (rows are gene names, columsn are sample identifiers) and produce a tab-delimited file as output.

### Signature matrices

The signature-matrices module implements four signature matrices − 2 derived from transcriptomics and two derived from proteomics measurements.

The mRNA-derived matrices are called LM22 and PBMC. The LM22 matrix was originally published in the CIBERSORT manuscript^14^ and contains expression values derived from microarray data for a group of filtered genes across 22 immune cell types and subtypes. The second published matrix PBMC (peripheral blood mononuclear cells) was derived from single-cell RNA sequencing (3’ sequencing) data in the CIBERSORTx m manuscript^19^ and comprises 8 immune phenotypes.

We also generated two additional signature matrices from a published proteomic dataset of flow cytometry-sorted PBMC^30^. Briefly, 28 distinct human hematopoietic cell types and subtypes from peripheral blood of healthy donors were sorted by flow cytometry. Erythrocytes and platelets were excluded from subsequent analyses. Cellular proteomes were analyzed in single runs by high-resolution MS using a quadrupole Orbitrap instrument. Each cell phenotype proteome was measured from four donors. The proteomic dataset included 10,134 proteins and 104 steady state samples. For LM9 we grouped the 26 phenotypes into 9 cell types: B cells, basophils, CD4 T cells, CD8 T cells, dendritic cells, eosinophils, monocytes, natural killer cells (NKs), neutrophils. For LM7c, basophils, eosinophils and neutrophils were grouped together as granulocytes. We took imputed values from Table S3 of the Rieckmann et al. paper^30^ to generate the two signature matrices, with samples first scaled to have zero mean and unit variance for LM9, using CIBERSORTx^19^ with these parameters: kappa= 999; q-value= 0.01; number of barcode genes= 300 to 500; disable quantile normalization= TRUE; filter non-hematopoietic genes= TRUE.

### Common Workflow Language deconvolution pipeline

We used the Common Workflow Language (CWL), following the syntax specified in CWL v1.2^37^, to link the individual docker images described above. Separate CWL script files were written for each step of data downloading, analyzing, and visualization. These individual script files have been integrated into ordered workflow steps in a single workflow file. Workflow has been primarily tested by the program cwltool, which is the reference implementation of programs that run CWL scripts, though can be employed using other CWL execution engines. The order of workflow steps was determined by using dependencies between the output of each step (e.g. data produced, file generated) and the input for the next step. The “scatter feature” was applied to facilitate parallel execution and accelerate the evaluations in each step. Essential results and log data were saved in order to retrieve or reanalyze the intermediate output files. The specification file for the workflow pipeline is written in YAML Ain’t Markup Language (YAML). The YAML files specify the input other parameters and/or arguments necessary for the pipeline.

### Docker image building

Each CWL file leverages a local Docker runtime to execute the underlying algorithm scripts. All individual steps are built into separate Docker images, which makes the pipeline reproducible and resolves the complexity of package management or issues arising from differing operating systems. The Docker images required to run the pipeline are included in the public image repository Docker Hub, at https://hub.docker.com/u/tumordeconv. When executed, each CWL performs a “pull action” and automatically downloads or updates the specified image it requires to complete its task. Docker images were automatically built with each code commit and pushed to the Github repository, using continuous integration and continuous deployment practices (CI/CD), to avoid conflicts that can arise with manually built or outdated images. Each commit push triggered a series of end-to-end tests on the CI/CD platform CircleCI, where the entire workflow is executed on a virtual machine. If the tests were successful, indicating the pipeline integrity was maintained with each code change, any associated Docker images were rebuilt and published to the repository.

### Data simulation

Pseudo-bulk data was simulated in a similar fashion as in Petralia et al (2022)^38^. Our simulation framework relied on two published datasets. First, we considered proteomic profiling from Rieckmann et al^30^. This study includes proteomic profiling of 26 immune cell subtypes, and then collapsed to k=9 different cell types: Neutrophils, Eosinophils, Basophils, B cells, CD4 T cells, CD8 t cells, Monocytes and Dendritic Cells. For each cell type *k*, 4 different proteomic profiles were provided, i.e., *μ*_1,*k*_, *μ*_2,*k*_, *μ*_3,*k*_, *μ*_4,*k*_. For each sample *i*, weights of different immune cells were randomly sampled from a dirichlet distribution with parameter 0.5 (i.e., *π*_*i*,1_, *π*_*i*,2_.. *π*_*i,K*_). Then, for each patient, mixed proteomic profiling was derived as the weighted average of proteomic profiling of different celltypes as follows:

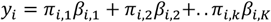

with *β*_*i,k*_ being one of the proteomic profiles available for the k-th cell type which was randomly sampled from *μ*_1,*k*_, *μ*_2,*k*_, *μ*_3,*k*_, *μ*_4,*k*_. Next, we considered data from Linsley et al.^39^, which contains transcriptomic profiling of 6 immune cell types including B-cells, CD4 t-cells, CD8 t-cells, Monocytes, Neutrophils and Natural Killers. For each cell type, this data contained 20 different transcriptomic profiling. Mixed transcriptomic data was generated similarly to proteomic profiling.

### Algorithm metrics

We use two types of metrics for comparing the deconvoluted results to *s* either simulated data or mRNA data from the same sample: namely a correlation-based metric and a distance based metric. The deconvoluted results are in a matrix where columns are the samples and rows are the cell type proportion calculated from the deconvolution algorithms. To compare any two deconvoluted matrices, we can calculate either the correlation or the distance between the corresponding vectors of cell type proportions. Given any two matrices *A* and *B* we can get cell type proportions *a*_\**j*_ ={*a*_1*j*_,…, *a*_*ij*_,…, *a*_*Nj*_} and *b*_\**j*_ ={*b*_1*j*_,…, *b*_*ij*_,…, *b*_*Nj*_} for patient *j*, where *N* is the number of cell types, and distributions across all patients *a*_*i*\*_ ={*a*_*i*1_,…, *a*_*ij*_,…, *a*_*i*,_} and *b*_*i*\*_ ={*b*_*i*1_,…, *b*_*ij*_,…, *b*_*i*,_} for cell type *i*, where *M* is the number of patients. We can then calculate the correlation and distances in the following approach.

### Correlation based comparison

In this comparison, each of the deconvoluted results are compared by calculating the Pearson correlation or Spearman correlation for each sample or for each cell type. The average correlation was simply calculated by averaging the correlation values across patients. The Pearson correlation for cell type proportions is calculated following:

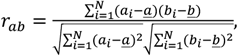

and the Spearman correlation for cell type proportions is calculated following:

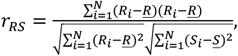

where, *R*_*i*_ and *S*_*i*_ are ranks of *a*_*i*_ and *b*_*i*_. For correlations between patients distributions, we replace the *N* with *M* in the equations above.

### Distance based comparison

In this comparison, we provide three different distance metrics, namely Euclidean, Jenson-Shannon divergence, Kolmogorov-Smirnov distance. For the distance metrics, we only calculate the distances between cell type proportions for each patient. An average distance was simply calculated by averaging the distance values across patients. The Euclidean distance is calculated following:

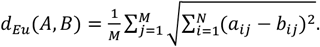

The Jenson-Shannon distance is calculated with:

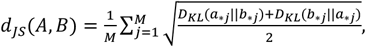

where *D*_*K*=(_*a*_\**j*_‖*b*_\**j*_) and *D*_*K*=(_*b*_\**j*_‖*a*_\**j*_) are the Kullback-Leibler (KL) divergences calculated by:

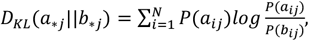

and *P*(*a*_*ij*_) is the proportion of cell type *i* in patient sample *j* in deconvoluted matrix *A* and similarity for *P*(*b*_*ij*_) in deconvoluted matrix *B*. For the Kolmogorov-Simirnov (KS) distance, we calculated the KS distance with the following equation:

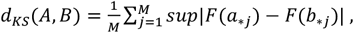

where *F*(*a*_\**j*_) and *F*(*b*_\**j*_) are the cumulative distribution function of *a*_\**j*_ and *b*_\**j*_.

## References

1. Fridman, W. H., Zitvogel, L., Sautés-Fridman, C. & Kroemer, G. The im-mune contexture in cancer prognosis and treatment. Nat. Rev. Clin. Oncol. 14, 717–734 (2017).

2. Hanahan, D. & Weinberg, R. A. Hallmarks of cancer: the next genera-tion. Cell 144, 646–74 (2011).

3. Baghban, R. et al. Tumor microenvironment complexity and thera-peutic implications at a glance. Cell Commun. Signal. 18, 59 (2020).

4. Gun, S. Y., Lee, S. W. L., Sieow, J. L. & Wong, S. C. Targeting immune cells for cancer therapy. Redox Biol. 25, 101174 (2019).

5. Ali, H. R. et al. Imaging mass cytometry and multiplatform genomics define the phenogenomic landscape of breast cancer. Nat. Cancer 1, 163–175 (2020).

6. Bodenmiller, B. Multiplexed Epitope-Based Tissue Imaging for Dis-covery and Healthcare Applications. Cell Syst. 2, 225–238 (2016).

7. Lun, X.-K. et al. Analysis of the Human Kinome and Phosphatome by Mass Cytometry Reveals Overexpression-Induced Effects on Cancer-Related Signaling. Mol. Cell 74, 1086-1102.e5 (2019).

8. Simoni, Y., Chng, M. H. Y., Li, S., Fehlings, M. & Newell, E. W. Mass cy-tometry: a powerful tool for dissecting the immune landscape. Curr. Opin. Immunol. 51, 187–196 (2018).

9. Pliner, H. A., Shendure, J. & Trapnell, C. Supervised classification enables rapid annotation of cell atlases. Nat. Methods 16, 983–986 (2019).

10. Liu, X. et al. Knowledge-based classification of fine-grained immune cell types in single-cell RNA-Seq data. Brief. Bioinform. 22, bbab039 (2021).

11. Azizi, E. et al. Single-Cell Map of Diverse Immune Phenotypes in the Breast Tumor Microenvironment. Cell (2018) doi:10.1016/j.cell.2018.05.060.

12. Sokolowski, D. J. et al. Single-cell mapper (scMappR): using scRNA-seq to infer the cell-type specificities of differentially expressed genes. NAR Genomics Bioinforma. 3, lqab011 (2021).

13. Finotello, F. & Trajanoski, Z. Quantifying tumor-infiltrating immune cells from transcriptomics data. Cancer Immunol. Immunother. CII 67, 1031–1040 (2018).

14. Chen, B., Khodadoust, M. S., Liu, C. L., Newman, A. M. & Alizadeh, A. A. Profiling tumor infiltrating immune cells with CIBERSORT. Methods Mol. Biol. Clifton NJ 1711, 243–259 (2018).

15. Aran, D., Hu, Z. & Butte, A. J. xCell: digitally portraying the tissue cellular heterogeneity landscape. Genome Biol. 18, 220 (2017).

16. Racle, J. & Gfeller, D. EPIC: A Tool to Estimate the Proportions of Dif-ferent Cell Types from Bulk Gene Expression Data. in Bioinformatics for Cancer Immunotherapy: Methods and Protocols (ed. Boegel, S.) 233–248 (Springer US, 2020). doi:10.1007/978-1-0716-0327-7_17.

17. Becht, E. et al. Estimating the population abundance of tissue-infiltrating immune and stromal cell populations using gene expression. Genome Biol. 17, 218 (2016).

18. Liu, C. C., Steen, C. B. & Newman, A. M. Computational approaches for characterizing the tumor immune microenvironment. Immunology 158, 70–84 (2019).

19. Newman, A. M. et al. Determining cell type abundance and expression from bulk tissues with digital cytometry. Nat. Biotechnol. 37, 773–782 (2019).

20. Racle, J., de Jonge, K., Baumgaertner, P., Speiser, D. E. & Gfeller, D. Simultaneous enumeration of cancer and immune cell types from bulk tumor gene expression data. eLife 6, e26476.

21. Sturm, G., Finotello, F. & List, M. Immunedeconv: An R Package for Unified Access to Computational Methods for Estimating Immune Cell Fractions from Bulk RNA-Sequencing Data. in Bioinformatics for Cancer Immunotherapy: Methods and Protocols (ed. Boegel, S.) 223–232 (Springer US, 2020). doi:10.1007/978-1-0716-0327-7_16.

22. Clark, D. J. et al. Integrated Proteogenomic Characterization of Clear Cell Renal Cell Carcinoma. Cell 179, 964-983.e31 (2019).

23. Dou, Y. et al. Proteogenomic Characterization of Endometrial Carcnoma. Cell 180, 729-748.e26 (2020).

24. Gillette, M. A. et al. Proteogenomic Characterization Reveals Therapeutic Vulnerabilities in Lung Adenocarcinoma. Cell 182, 200-225.e35 (2020).

25. Zhang, H. et al. Integrated proteogenomic characterization of human high grade serous ovarian cancer. Cell 166, 755–765 (2016).

26. Fortelny, N., Overall, C. M., Pavlidis, P. & Freue, G. V. C. Can we predict protein from mRNA levels? Nature 547, E19–E20 (2017).

27. McManus, J., Cheng, Z. & Vogel, C. Next-generation analysis of gene ex-pression regulation--comparing the roles of synthesis and degrada-tion. Mol. Biosyst. 11, 2680–2689 (2015).

28. Payne, S. H. The utility of protein and mRNA correlation. Trends Bio-chem. Sci. 40, 1–3 (2015).

29. Nagaraj, N. et al. Deep proteome and transcriptome mapping of a hu-man cancer cell line. Mol. Syst. Biol. 7, 548 (2011).

30. Rieckmann, J. C. et al. Social network architecture of human immune cells unveiled by quantitative proteomics. Nat. Immunol. 18, 583–593 (2017).

31. Avila Cobos, F., Alquicira-Hernandez, J., Powell, J. E., Mestdagh, P. & De Preter, K. Benchmarking of cell type deconvolution pipelines for tran-scriptomics data. Nat. Commun. 11, 5650 (2020).

32. Decamps, C. et al. DECONbench: a benchmarking platform dedicated to deconvolution methods for tumor heterogeneity quantification. BMC Bioinformatics 22, 473 (2021).

33. Jin, H. & Liu, Z. A benchmark for RNA-seq deconvolution analysis un-der dynamic testing environments. Genome Biol. 22, 102 (2021).

34. Thorsson, V. et al. The Immune Landscape of Cancer. Immunity 48, 812-830.e14 (2018).

35. Gibbs, D. L. Robust classification of Immune Subtypes in Cancer. 2020.01.17.910950 Preprint at https://doi.org/10.1101/2020.01.17.910950 (2020).

36. Lindgren, C. M. et al. Simplified and Unified Access to Cancer Proteo-genomic Data. J. Proteome Res. 20, 1902–1910 (2021).

37. Amstutz, P. et al. Common Workflow Language, v1.0. (2016) doi:10.6084/m9.figshare.3115156.v2.

38. Petralia, F. et al. BayesDeBulk: A Flexible Bayesian Algorithm for the Deconvolution of Bulk Tumor Data. 2021.06.25.449763 Preprint at https://doi.org/10.1101/2021.06.25.449763 (2022).

39. Linsley, P. S., Speake, C., Whalen, E. & Chaussabel, D. Copy number loss of the interferon gene cluster in melanomas is linked to reduced T cell infiltrate and poor patient prognosis. PloS One 9, e109760 (2014).

